# Prioritization of The Zinc finger domain within *BCL11A by the* Amelioration capability of hemoglobinopathies using CRISPR-Cas9 technology

**DOI:** 10.1101/2023.11.28.568974

**Authors:** Kainaat Mumtaz, Irfan Hussain, Fizza Iftikhar, Fawad U Rehman, Muhammad Jameel, Safana Farooq, Hammad Hassan, Ambrin Fatima, Oliver Gerhard Ottmann, Afsar Ali Mian

**Affiliations:** Centre for Regenerative Medicine and Stem Cells Research, The Aga Khan University, 1st Flour,Juma Building, Stadium Road, Karachi 74800, Sindh, Pakistan; Department of Hematology, Cardiff University, Cardiff, United Kingdom

**Keywords:** sickle cell disease, Thalassemia, BCL11A gene, Fetal Hemoglobin, Crispr cas9

## Abstract

BCL11A/EVI9, a zinc-finger protein primarily expressed in brain and hematopoietic cells, plays a central role in lymphocyte development, gamma-globin suppression, spinal neuron development, sensory innervation, neuronal polarity, migration, and is associated with microcephaly and dysregulated brain-related genes, offering therapeutic potential for sickle cell disease. The function of the transcriptional regulator is intricately linked to its structural organization, which determines its ability to interact with specific DNA sequences and modulate gene expression. BCL11A boasts multiple domains, including six C2H2 zinc fingers, a C2HC zinc finger, a NuRD-interacting domain, an acidic domain, and a proline-rich domain. In the present study, we delve into the intricate structure and function of the zinc finger domains located in the BCL11A gene, which plays a crucial role in regulating the expression of gamma-globin gene. Specifically, three C2H2-type zinc finger domains, Znf4, Znf5, and Znf6, within BCL11A, are known to bind to DNA. Znf4 and Znf5 demonstrate a significant interaction with the TGACCA motif in the gamma-globin −115 HPFH region sequence, contributing substantially to DNA binding specificity. Although Znf3 and Znf6 also interact with DNA, their contributions are comparatively minor. Employing CRISPR-Cas9 technology, targeted genomic deletions of Znf4 exhibit high efficiency, opening doors for further research. Edited CD34+ cells successfully differentiate into erythrocytes without impairments, underscoring CRISPR-Cas9’s suitability for studying gene functions in erythropoiesis. Furthermore, BCL11A knockdown via sgRNAs results in elevated gamma-globin expression, offering a promising therapeutic avenue for beta-hemoglobinopathies. HPLC analysis reveals a substantial increase in HbF levels, particularly upon Znf4 deletion, emphasizing BCL11A gene potential as a therapeutic target. These findings also highlight the connection between the function of BCL11A and its structural organization, which can be modulated, and this insight can potentially be extended to uncover its roles in various other domains.

## Introduction

The phenotypic heterogeneity of the hemoglobin disorders contrasts with their Mendelian inheritance(Angastiniotis, Vives Corrons, Soteriades, & Eleftheriou, 2013; Modell & Darlison, 2008). The clinical severity of the β-hemoglobin disorders is primarily determined by the levels of fetal hemoglobin (HbF, α2γ2)(Amaya et al., 2013; Antoniani et al., 2018). In β-thalassemia, increased levels of γ-globin substitute for absent β-globin, thus reducing relative α-chain excess(Bauer et al., 2013). In sickle cell disease (SCD), γ-globin acts as a potent anti-sickling agent(Borg et al., 2010). Clinical observations, including those of rare patients co-inheriting highly elevated HbF (hereditary persistence of fetal hemoglobin), large cohorts with varying HbF levels, and the natural history of infants with waning HbF, unequivocally establish that HbF mitigates the β-hemoglobin disorders(Breda et al., 2016; Chang et al., 2017; Costa, Capuano, Sommese, & Napoli, 2015; Galarneau et al., 2010).

An understanding of the mechanisms controlling hemoglobin switching, particularly the fetal-to-adult transition, was once elusive despite extensive inquiry(Huang et al., 2017; Ivaldi et al., 2018). This stems from the challenge of imposing stage specificity for globin transcription in an erythroid cell context, where the shift in transcript production from the γ-genes to the β-gene in the β-like globin complex occurs(Krasilnikova & Karamjan, 2017). However, uncovering these mechanisms is crucial as the reactivation of HbF has the potential to cure β-hemoglobin disorders(Sankaran et al., 2011). A therapy that results in measurable amounts of HbF in all erythrocytes, rather than limited to a subset, is optimal(Martyn et al., 2018). Therefore, understanding the molecular circuitry controlling HbF is essential for rationally designed therapies(Shi, Cui, Engel, & Tanabe, 2013).

Recent research has identified BCL11A as a critical modifier of hemoglobin disorders(Basak et al., 2015). These findings have direct therapeutic relevance for the hemoglobinopathies and emphasize the importance of contextual gene regulation in determining heritable disease severity(Basak et al., 2020). The zinc-finger transcriptional factor BCL11A plays a crucial role in regulating fetal hemoglobin (HbF) switching by silencing embryonic and fetal globin genes during development(Bauer et al., 2013; Chen, Luo, Steinberg, & Chui, 2009). In both mouse models and human cells, BCL11A has been shown to associate with known γ-globin transcriptional repressors and bind to the locus control region and other intergenic sites, which prevents the interaction between the locus control region and the HbF globin gene required for fetal globin expression(Canver et al., 2015). Studies have demonstrated that individuals with incomplete hemoglobin switching and elevated HbF levels in adult life exhibit milder symptoms of sickle cell disease and β-thalassemia(Ippolito et al., 2014). Genome-wide association studies have also revealed that variations in HbF levels are frequently caused by common polymorphisms in the BCL11A gene and the β-globin cluster(Canver et al., 2015; Sankaran et al., 2009; Satterwhite et al., 2001).

The critical role of BCL11A in repressing γ-globin gene expression is well-supported by genetic evidence, however, the exact mechanism and location of its action have been a topic of debate(Xu et al., 2010). Conflicting reports exist regarding whether BCL11A or its paralog BCL11B directly binds DNA, and if so, which sequence(s) it recognizes(Uda et al., 2008). Previous studies have suggested that BCL11A acts at a distance from the γ-globin gene promoters, but the exact sites of occupancy within the β-globin locus have not been precisely mapped(Smith et al., 2019). To gain a better understanding of BCL11A’s mode of action, researchers have focused on identifying its essential domains and DNA binding specificity at the HBF locus(Martyn et al., 2018). In the present study, we delve into the intricate structure and function of the zinc finger domains located in the BCL11A gene, which plays a crucial role in regulating the expression of gamma-globin gene. Our research employs the powerful tool of CRISPR-Cas9-mediated genome editing to investigate the function of these domains in biological processes, as well as to explore the potential for therapeutic strategies targeting this gene.

## Methodology

### BCL11A and −115 HPFH Region of Gamma-Globin Gene interactions analysis

In 2020, Dr. Xu Liu and a team of researchers from the University of California, San Francisco, published a study in the scientific journal Nature, outlining the crystal structure of BCL11A in conjunction with the −115 HPFH region of the gamma-globin gene(Yang et al., 2019). The PDB structure 6KI6 was analyzed using the DNAproDB (https://dnaprodb.usc.edu) which a web-based database and structural analysis tool (Sagendorf, Markarian, Berman, & Rohs, 2020).

### Cell line culturing

To sustain the growth of K562 cell line, a nourishing RPMI 1640 medium from Lonza was utilized. This medium was enhanced with crucial nutrients, such as 10% fetal bovine serum and glutamine, both of which were also obtained from Lonza. The addition of Hepes from Life Technologies maintained the optimal pH levels, while sodium pyruvate from the same source provided the necessary energy levels. Furthermore, penicillin and streptomycin from Life Technologies were added to the medium to prevent bacterial contamination and ensure a sterile cell culture environment.

### Isolation, characterization, and differentiation of CD34+

CD34+-enriched cells derived from peripheral blood of both healthy and patient. CD34+ cells were isolated using Lymphoprep density gradient centrifugation, followed by positive selection with anti-CD34-tagged magnetic beads using MACS LS columns. The CD34+ cells were then cultured under different conditions, including proliferation, expansion, and differentiation, for a period of 21 days. To achieve this, the cells were first expanded with stem cell factor (SCF), FLT-3, and TPO, along with dexamethasone, until day 7 of culture. Subsequently, the cells were cultured in a first differentiation media supplemented with erythropoietin (EPO) from day 7 to 14 to observe early erythroblast cells. Finally, the cells were transferred to a second differentiation media containing EPO and holo-transferrin until the 21st day of culture to observe terminally differentiated erythroid cells. The cells were characterized at different time points using flow cytometry. The detailed protocol is given in the supplementary file.

### gRNA design and Transfection

We employed the CRISPOR (45) online tool to design gRNAs targeting znf2 and znf4 within exon 2 and exon 4 of the BCL11A gene, respectively. The resulting oligonucleotide and the recombinant Cas9 protein fused with EGFP, the custom-designed crRNA, and tracrRNA were procured from Integrated DNA Technologies in Coralville. The sequences of these components are available in the supplementary table. The individual components were combined to form the RNP complex, which was used in accordance with the transfection protocol provided on IDT’s website for the Alt-R CRISPR/Cas9 System. The Neon Transfection parameters, including voltage (1,150–1,700 V), duration (10–40 ms), and pulse (1–3), were optimized to achieve maximum efficiency of RNP delivery into CD34+ cells. The ideal conditions for RNP delivery were found to be 1,600 V, 10 ms, and 3 pulses. The transfected CD34+ cells were transferred into 6-cell plates with 2 mL of pre-warmed, complete CD34+ cell culture medium, and grown in culture for 16–72 hours. Subsequently, the cells were harvested and subjected to flow cytometry to assess the delivery efficiency of the EGFP RNP complex.

### Differentiation of CD34+ cells into erythrocytes

CD34+ cells were cultured in a two-stage differentiation process using distinct media formulations. During the initial stage, cells were cultured in differentiation media supplemented with erythropoietin (EPO) from day 7 until day 14 to observe the emergence of early erythroblast cells. Subsequently, the cells were transferred to a second differentiation media containing both EPO and holo-transferrin and cultured until day 21 to observe terminally differentiated erythroid cells. Flow cytometry analysis revealed that the highest recovery of CD34+ cells occurred on day 3 of culture under all conditions. The majority of cells were found to be pro-erythroblasts at day 7, with maturation progressing to mature erythroblasts by day 14-21. The presence of transferrin and EPO favored erythroid differentiation and maturation and decreased the proportion of non-erythroid CD45+ cells (Figure 4). Overall, this culture method and condition produced a substantial quantity of pure erythroid cells from a small amount of peripheral blood, providing a reliable in vitro model of human erythropoiesis.

### Assessment of genome editing outcomes

The genome editing events were examined in K562 cells at the onset and after 9 days of erythroid differentiation, as well as in erythroid cells derived from adult mobilized HSPCs after 6 and 14 days of erythroid differentiation, correspondingly. Genomic DNA from both control and edited cells was extracted using the PureLink Genomic DNA Mini Kit (Life Technologies), Quick-DNA/RNA Miniprep (ZYMO Research), or DNA Extract All Reagents Kit (Thermo Fisher Scientific), following the respective manufacturer’s guidelines. To assess the efficacy of NHEJ at gRNA target sites, we carried out PCR followed by Sanger sequencing and TIDE analysis (tracking of InDels by decomposition) (49) or ICE CRISPR Analysis Tool (Synthego)

### RT-qPCR analysis of globin and erythroid markers

The RNeasy Micro kit (Qiagen) was used to extract total RNA from k562, CD34+ and primary mature erythroblasts, according to the manufacturer’s instructions. SuperScript First-Strand Synthesis System for RT-qPCR (Invitrogen) with oligo(dT) primers was used to reverse transcribe mature transcripts. RT-qPCR was carried out with an iTaq Universal SYBR Green master mix (Bio-Rad).

### Western blotting

The western blot was conducted according to the method described earlier (Mahmood & Yang, 2012). In brief, samples were treated with 1x SDS loading buffer to denature proteins and subjected to separation by 13% or gradient SDS-PAGE gels. The separated proteins were then transferred onto a PVDF membrane using a standard wet transfer system at 2.5 mA/cm2 for 2 hours. The membrane was blocked with 5% nonfat milk for 1 hour and then incubated with primary antibodies for 1 hour at room temperature or overnight with shaking at a cold room. The excess antibodies were washed with TBS-T (50 mM Tris pH 8.0, 150 mM NaCl, 0.1% Tween 20) for three times, and then HRP-conjugated secondary antibodies were added and incubated for 30 minutes at room temperature. After washing the membranes three times with TBS-T, they were developed using Immobilon Western Chemiluminescent HRP Substrate (Millipore, WBKLS0500). The primary antibodies used were M2-Flag (F1804, Sigma-Aldrich), BCL11A (ab19487 and ab191401, Abcam), and GAPDH (sc-25778, Santa Cruz Biotechnology), all diluted at 1:1,000 in TBS-T.

### Hb quantification by HPLC

The HPLC analysis was conducted using the LC Solution software (Shimadzu) and a NexeraX2 SIL-30 AC chromatograph. Hb tetramers were separated using two cation-exchange columns (PolyCAT A, PolyLC, Columbia, MD) with elution carried out by a gradient mixture of solution A [20 mM Bis-Tris and 2 mM KCN (pH 6.5)] and solution B [20 mM Bis-Tris, 2 mM KCN, and 250 mM NaCl (pH 6.8)]. The absorbance at 415 nm was measured to determine the total Hb levels. The calculation of Hb levels was based on the integration of the areas under the Hb peaks and comparison with a standard Hb control (Lyphochek Hemoglobin A2 Control, Bio-Rad).

### Statistical analysis

The statistical analysis was carried out using R programming. Paired t-tests were performed to compare genome editing efficiencies in erythroid subpopulations sorted based on HbF expression, while unpaired t-tests were used for all other analyses. The Kruskal-Wallis test was used to compare the frequency of deletion generated at each nucleotide by the different gRNAs. The statistical significance was set at a threshold of P < 0.05.

## Results

### The Structure and Function of Zinc Finger Domains in BCL11A: Implications for the Regulation of Gamma-Globin Gene Expression

BCL11A is a gene that comprises three C2H2-type zinc finger domains: Znf4, Znf5, and Znf6. These domains are located in respective exons and are known to bind to DNA. The first two zinc finger domains of BCL11A are highly conserved among different species, indicating their crucial role in regulating gene expression. In contrast, the third zinc finger domain exhibits more variation and may have a role in interacting with other proteins. The zinc finger domains of BCL11A, specifically Znf4 and Znf5, adopt the canonical C2H2 zinc finger fold. These domains wrap around the DNA duplex and interact with a total of 7 bp (−119)(TTGACCA)(−113) from the γ-globin −115 HPFH region sequence. While Znf4 interacts with the 4-bp TTGA sequence, Znf5 interacts with all 6 bp of the TGACCA motif. In the BCL11A structure, Znf4-5 makes numerous hydrogen-bonding and van der Waals contacts with both DNA strands to contribute to the binding affinity and specificity. Although Znf3 and Znf6 also interact with the DNA duplex, they contribute less significantly.

### Investigating the Role of ZNF4 and Znf2 with CRISPR-Cas9-Mediated Targeted Genomic Deletions

The present study aimed to employ CRISPR-Cas9 technology with sgRNA pairs to generate targeted genomic deletions of ZNF4 exon in CD34 positive cells and K562 cells. RNP transfection was performed to induce genomic deletions, and transfection efficiency was assessed, which was approximately 70% for K562 cells and 54% for CD34 cells. The FACS system was utilized to sort K562 cells 24 hours post-electroporation, and they were plated at a limiting dilution. Subsequently, genomic DNA was isolated from the transfected cells, and deletion events were analyzed by PCR using primer pairs flanking the sgRNA recognition sites. The obtained PCR products confirmed a 200bp genomic deletion for all tested sgRNA combinations, and heterozygote, homozygote, and non-deleted clones were identified. Additionally, the PCR products were gel purified and subjected to direct Sanger sequencing. Notably, about 32% (8/25) of the single clones were identified as biallelic deleted clones. These findings indicate that CRISPR-Cas9-mediated genome editing is a highly effective approach to induce targeted genomic deletions of ZNF4, and the use of sgRNA pairs can further enhance the efficiency of this method. These results have important implications for exploring the role of ZNF4 in biological processes and developing potential therapeutic strategies targeting this gene.

### CRISPR-Cas9 edited CD34+ cells differentiate into erythrocytes without impairment

Next, we investigated the differentiation of CRISPR edited CD34+ cells into erythrocytes and monitor the expression of surface antigens, including CD71 and GPA, which are specific markers for cells of the erythroid lineage. To induce differentiation, cells were cultured in the presence of EPO, insulin, and holo-transferrin after transfection. Flow cytometry analysis was performed to determine the percentage of CD71+ and GPA+ cells. The results showed a significant increase in both CD71+ and GPA+ cells in the transfected and differentiated control group compared to undifferentiated CD34+ cells. Furthermore, edited cells exhibited an increase in CD71+ cells from 6.4% to 9.9% (1.54-fold increase) and a significant increase in CD235a+ cells from 4% to 6.6% (1.65-fold increase) on day 4 of differentiation, as depicted in Figure 10. These findings suggest that CRISPR-Cas9-mediated genome editing does not impair the differentiation potential of CD34+ cells into erythrocytes and may be utilized for future research on the role of specific genes in erythropoiesis.

### BCL11A knockdown induces gamma-globin expression: Evidence from qRT-PCR analysis and western blotting

Next, we aimed to evaluate the efficacy of BCL11A inhibition using sgRNAs, with a specific focus on reactivating fetal hemoglobin. To this end, we conducted a thorough analysis of BCL11A expression at the mRNA level in bulk and edited clones, utilizing quantitative real-time PCR (Fig. 2A). Furthermore, we investigated the effect of 200 bp deletions of Znf4 and Znf2 of BCL11A on gamma-globin expression by quantifying gamma-globin mRNA transcripts using quantitative RT-PCR (Figure). Intriguingly, our results revealed a significant increase in gamma-globin expression in both CD34 and K562 cells when targeting ZNF4, while the effect of Znf2, which leads to the loss of all zinc finger domains, was comparatively low on fetal hemoglobin induction.

**Figure 1.**
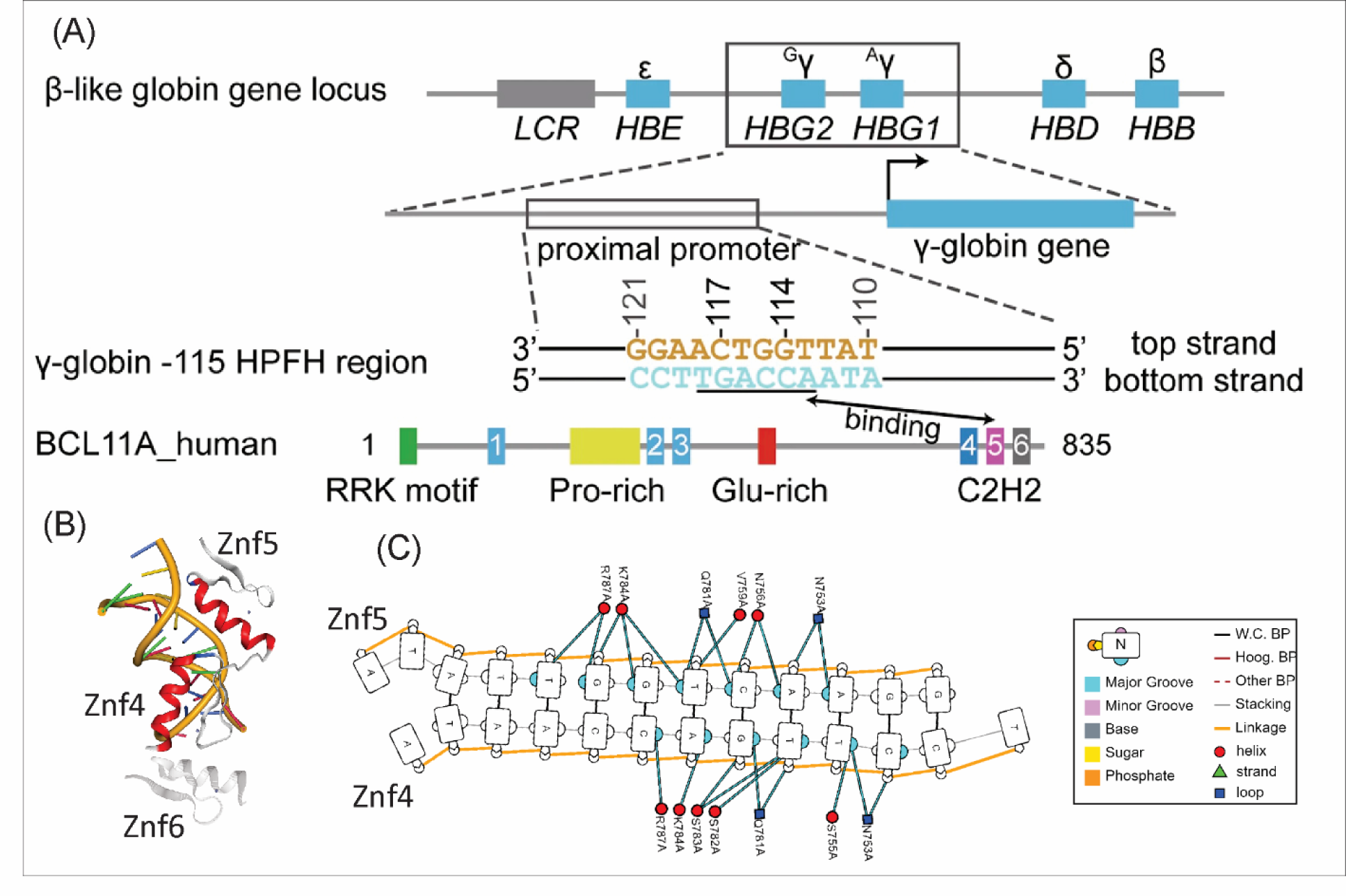
Examining the Relationship Between Znf4-5 of BCL11A and the γ-Globin −115 HPFH Region Sequence. (A) Upper panel: A visual overview of the β-like globin gene locus, focusing on the magnified depiction of the γ-globin genes and their adjacent promoters. In the middle panel, we present the structural representation of the γ-globin −115 HPFH region sequence, originating from the proximal promoter situated approximately 115 bp upstream of the transcription start sites for γ-globin genes. The strand containing the TGACCA sequence (marked by underlining) is denoted as the bottom strand, while its complementary counterpart is known as the top strand. These strands are distinguished by their orange and cyan colors, respectively. The two HPFH sites, G:C−117 and C:G−114, are numerically indicated above the base pairs. The lower panel offers an illustrative outline of the human BCL11A domain configuration, accentuating the interaction involving Znf4-6 of BCL11A and the γ-globin −115 HPFH region sequence. (b) The complete structure of Znf4-6 of BCL11A, forming a complex with the γ-globin −115 HPFH region sequence, is showcased in a simplified diagram. Znf4, Znf5, and Znf6 are color-coded (C) The Residue Contact Map shows individual nucleotide-residue interactions, DNA secondary structure, protein secondary structure and DNA interaction moieties. The DNA is displayed as a graph, with nucleotides being nodes and edges between them indicating backbone links, base pairing or base stacking. Different base pairing geometries are indicated via the base-pair edges, and other structural features such as backbone breaks, missing phosphates, and the DNA strand sense are represented. Protein residues are displayed as small nodes with the node shape and color representing residue secondary structur. Edges between residue and nucleotide nodes represent an interaction between the two and which DNA moiety(s) the interaction involves.

**Figure 2:**
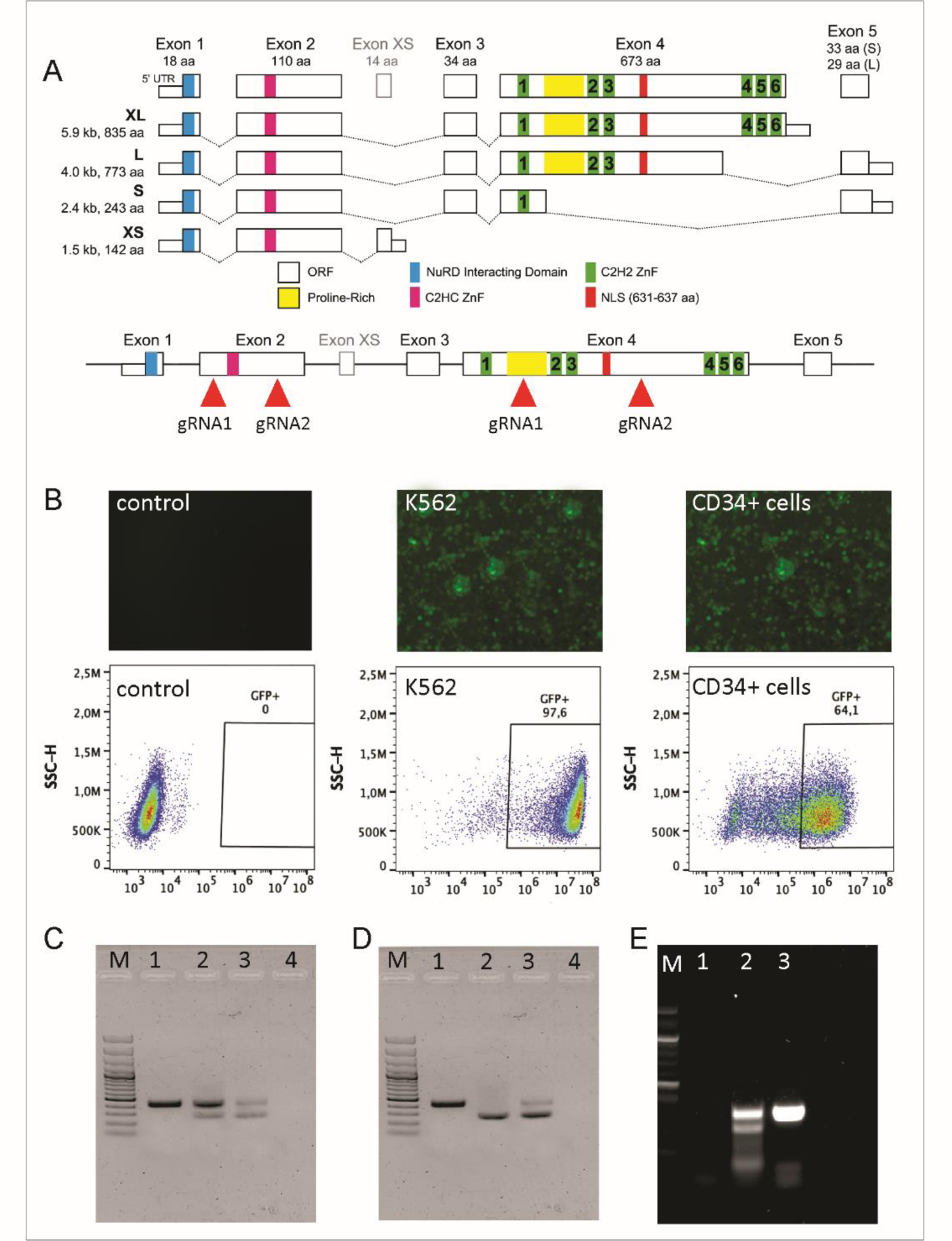
Comprehensive Analysis of Human BCL11A Isoforms: Deletion Validation and Expression Profiling in K562 and cd34+ Cells (A) The schematic diagram displays the various isoforms of human BCL11A. The red rectangle indicates the precise location of the guide RNA (gRNA). In Figure B, the expression of GFP is exhibited across different cell groups including control, K562 cells, and cd34+ cells. Furthermore, their fluorescence-activated cell sorting (FACS) analysis is presented. Moving to Figure C, a PCR analysis validates the deletions within the bulk population. Notably, the 100bp marker (M) is used for size reference. The K control amplification of 580 bp is represented by 1, whereas B2 and B3 illustrate both wild-type and deleted bands of 210 bp in K562 and cd34+ cells. Additionally, $4 signifies the non-template control. In Panel D, the validation of deletion within the pure population is demonstrated. This is also accompanied by the 100bp marker (M), control amplification (1) of 580 bp, and bands (2) and (3) representing wild-type and deleted sequences of 210 bp in K562 and cd34+ cells, along with the non-template control (4). The presence of a double band in K562 cells is attributed to the multiple copies of BCL11A. Figure E provides insight into the T7 assay. Here, the marker (M) is included, along with the non-template control (1), the verification of deletion through the T7 endonuclease assay (2), and the non-edited control (3).

The inhibition of BCL11A for reactivation of fetal hemoglobin expression represents a promising therapeutic avenue for the treatment of β-hemoglobinopathies. To assess the levels of fetal globin protein expression, Western blot analysis was employed on the three clones exhibiting the highest gamma-globin gene expression. Our findings demonstrated that the levels of fetal globin protein were consistent with the qRT-PCR results (Figure 23), providing additional evidence for the effectiveness of BCL11A knockdown in inducing gamma-globin expression. These results accentuate the potential of BCL11A knockdown as a therapeutic approach for the treatment of β-hemoglobinopathies and reinforce the need for further research to refine the clinical implementation of this strategy.

**Figure 3:**
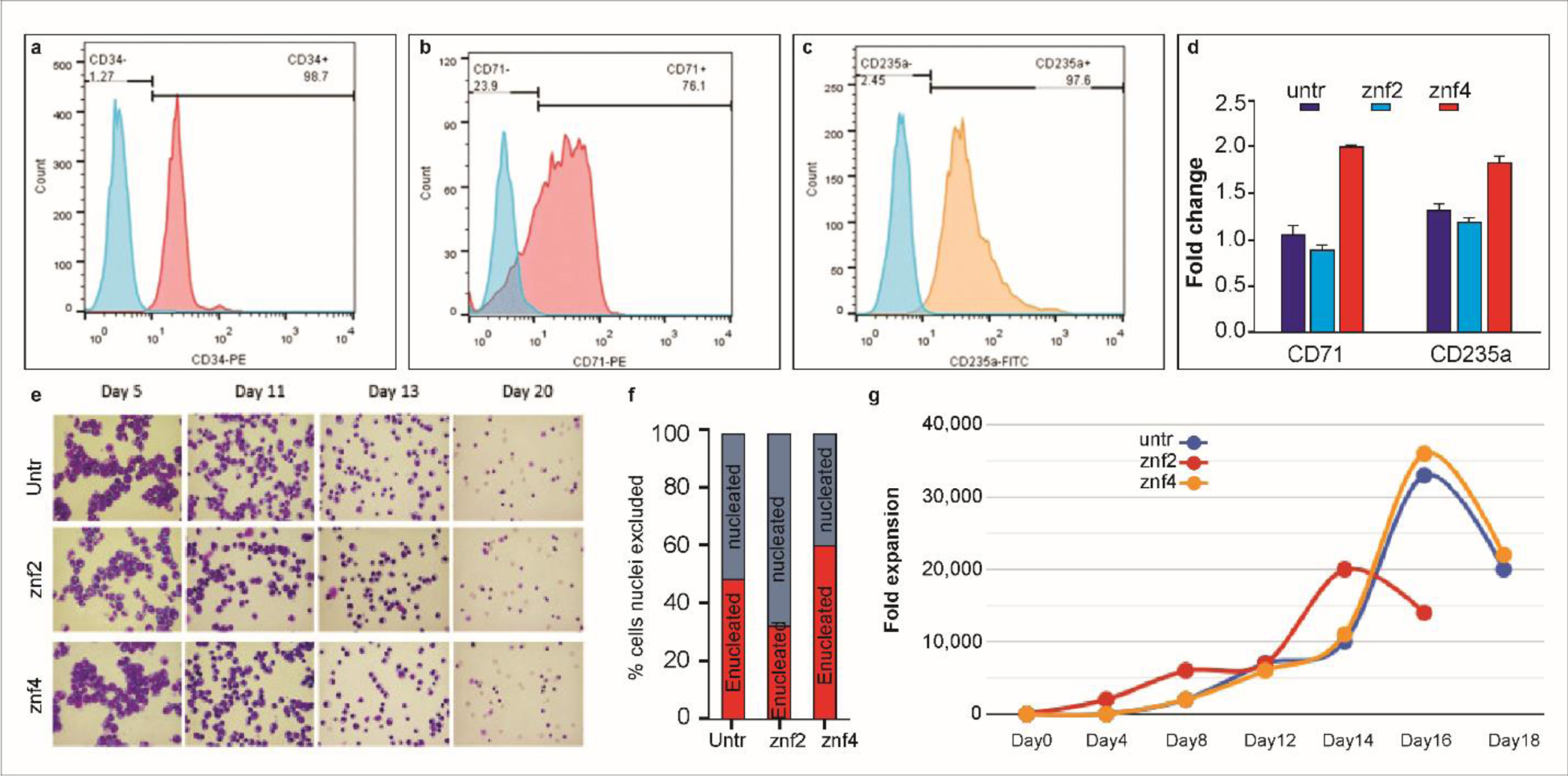
Characterization of CRISPR-Edited CD34+ Hematopoietic Stem Cells (HSCs) and Erythroid Precursors: The analysis involved the assessment of distinct markers, namely CD34 for HSCs, and CD71 and CD235a for erythroid precursors. In Figure a, a notable 98% of the isolated cells exhibited the expression of CD34. Additionally, Figures b and c demonstrate the presence of erythroid precursors marked by the expression of both CD71 and CD235a. Furthermore, in Panel D, it can be observed that CD34+ cells with BCL11A ZNF4 deletions showcased an augmented expression of surface markers associated with proerythroblastic and terminally mature erythrocytes. This heightened expression is in contrast to the untreated and znf2 deleted of edited CD34+ cells. (e) Hematoxylin and Eosin (H&E) Staining of Cytospin Preparations at Various Erythroid Differentiation Time Points.(f) Proportion of Nucleated to Enucleated Cells Assessed at the Culmination of the Culture.(g) Progression of Erythroid Differentiation in Untransfected (Untr), Exon 2 Zinc Finger Domain (znf2), and Exon 4 (znf4) Transfected Erythroid Cells, as Indicated by Growth Patterns.

**Figure 4:**
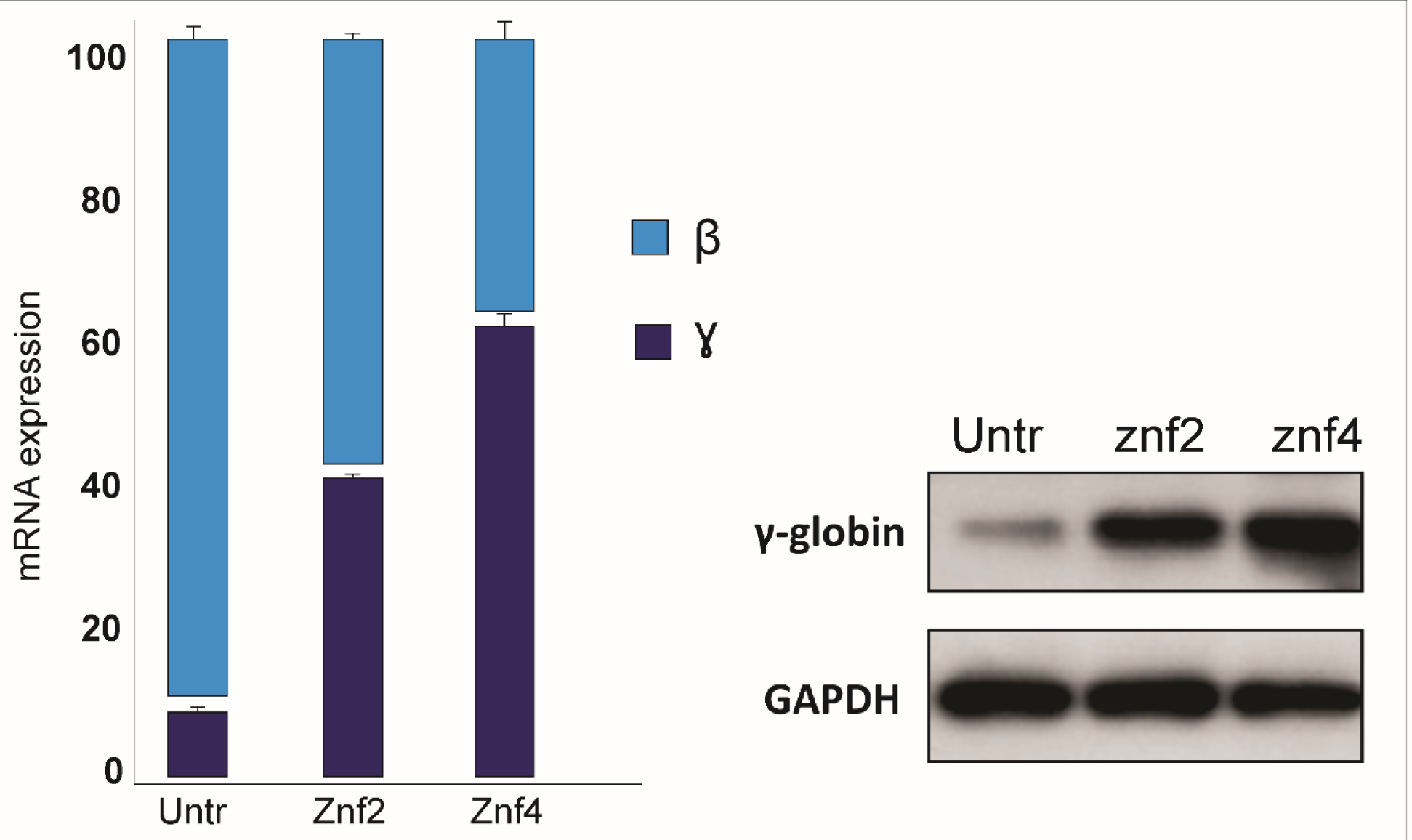
Quantitative PCR and western blot analysis for the confirmation of induction of fetal hemoglobin.

### Znf4 deletion boosts HbF expression in gene-edited cells: HPLC analysis

The protein-level HbF expression in gene-edited CD34+ HSPCs and K562 cells was examined using HPLC-mediated hemoglobin electrophoresis. Of note, comparative analysis was conducted between the results obtained from the deletion of both Znf2 and Znf4 domains. Remarkably, hemoglobin electrophoresis disclosed that all edited samples exhibited a substantial increase in HbF levels as compared to the control group (Figure 5). Notably, the induction observed in the edited samples, particularly with Znf4 deletion, was significantly higher than that observed in the control and Znf2-edited samples.

**Figure 5:**
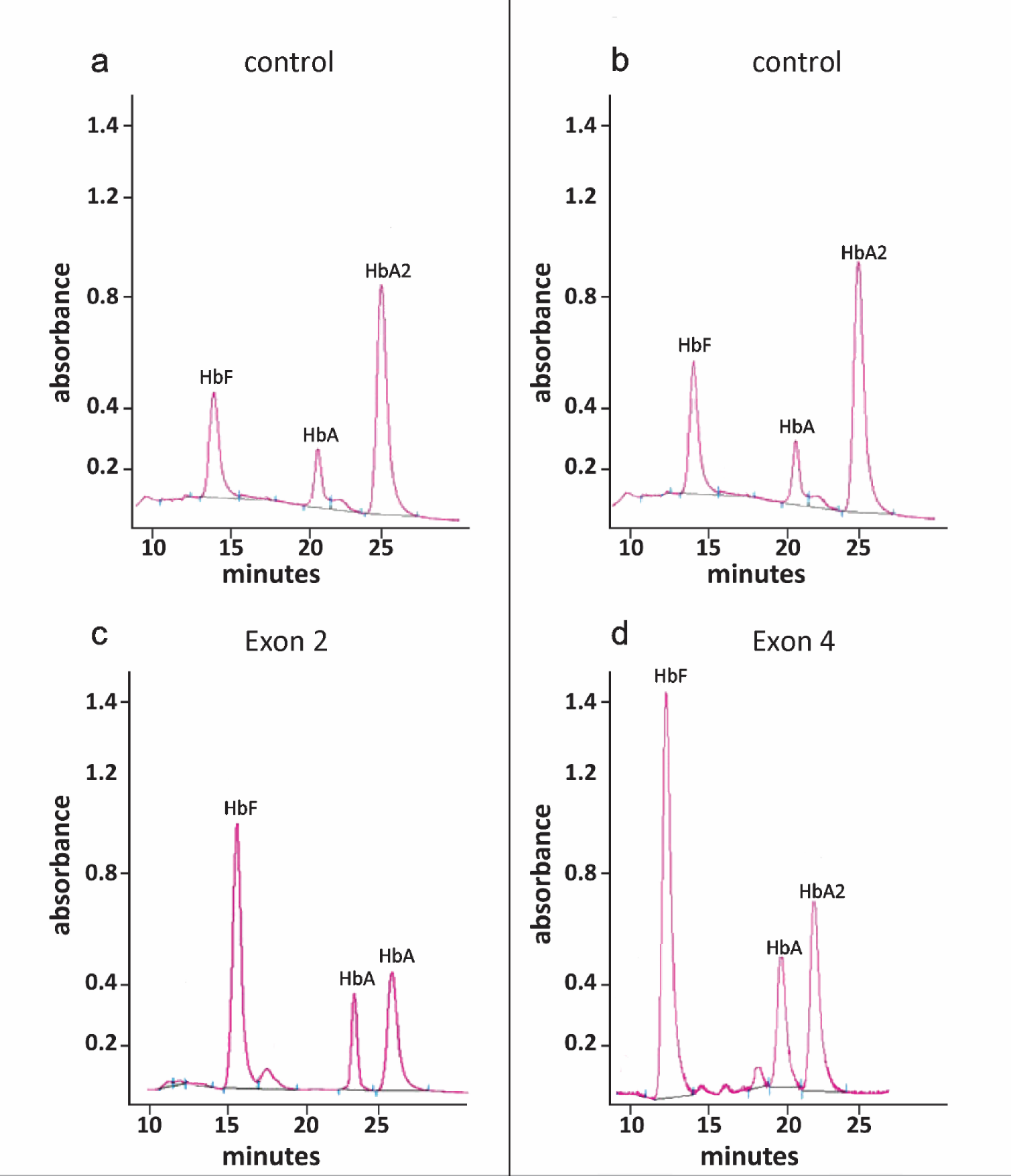
HbF expression in gene-edited CD34+ HSPCs cells examined using HPLC-mediated hemoglobin electrophoresis. Panel 1 is the control and exon1 deleted CD34+ HSPCs. While the second panel depicts the control and the Znf4 harboring exon 4 deleted CD34+ HSPCs.

## Discussion

BCL11A/EVI9, a zinc-finger protein primarily expressed in brain and hematopoietic cells, plays a central role in lymphocyte development, gamma-globin suppression, spinal neuron development, sensory innervation, neuronal polarity, migration, and is associated with microcephaly and dysregulated brain-related genes, offering therapeutic potential for sickle cell disease(Costa et al., 2015; Smith et al., 2019; Yang et al., 2019). The function of the transcriptional regulator is intricately linked to its structural organization, which determines its ability to interact with specific DNA sequences and modulate gene expression(Ippolito et al., 2014; Xu et al., 2010).

In a study by Liu et al. (2003), exon 1 of the Bcl11a gene was deleted in embryonic stem cells to create Bcl11a-deficient mice. While heterozygous mice remained viable and fertile, Bcl11a −/− mice had a brief lifespan of 3 to 4 hours after birth. Analysis of Bcl11a −/− fetal liver cells revealed the absence of B-lymphocyte markers and T-cell alpha/beta receptors but normal development in other cell lineages. Xu et al. (2011) demonstrated that the repressor BCL11A is essential for silencing gamma-globin expression in adult animals but not for red cell production. BCL11A acts as a barrier to HbF reactivation. Inactivation of BCL11A in sickle cell disease transgenic mice corrected disease-associated defects through high-level pancellular HbF induction, indicating BCL11A as a potential therapeutic target for sickle cell disease. John et al. (2012) conducted conditional knockout of Bcl11a in mouse spinal cord, revealing its role in the differentiation and morphogenesis of dorsal spinal neurons. Bcl11a was also necessary in spinal target neurons for proper wiring and differentiation in the superficial dorsal horn. Dysregulated Frzb expression contributed to sensory innervation disruptions in Bcl11a mutant spinal cord. Wiegreffe et al. (2015) used conditional knockout mice to establish Bcl11a’s requirement in neuronal cell polarity switching and migration of upper-layer projection neurons. Bcl11a −/− neurons exhibited altered morphology, orientation, and slowed migration, partly due to repression of the guidance cue Sema3c by Bcl11a. Dias et al. (2016) found that Bcl11a haploinsufficiency in mice resulted in microcephaly, brain volume reduction, abnormal behavior, and transcriptional deregulation in the cortex and hippocampus, affecting genes involved in ion transport, membrane trafficking, and neuronal signaling.

The gene’s functionality is determined by its various domains, with one such domain being the Zinc Finger family(Avram et al., 2000; Menzel et al., 2007; Wolfe, Nekludova, & Pabo, 2000). Zinc finger proteins (ZFPs), comprising the largest family of transcription factors characterized by finger-like DNA binding domains, exert a significant influence on diverse biological processes(Nakamura et al., 2000). In the context of tumorigenesis and tumor progression, ZFPs primarily function as transcription factors(Pedone et al., 1996). These proteins, known as transcription factors (TFs), assume a crucial role in intricate biological phenomena such as metabolism, autophagy, apoptosis, immune responses, stemness maintenance, and differentiation(Altshuler et al., 2010). TFs modulate gene transcription by directly recognizing or binding to DNA sequences(Frontini et al., 2002). The classification of zinc finger motifs reveals eight distinct categories based on their main-chain conformation and secondary structure surrounding their zinc-binding sites(Marban et al., 2005). These categories include Cys2His2 (C2H2)-like, Zn2/Cys6, Treble clef, Zinc ribbon, Gag knuckle, TAZ2 domain-like, Zinc binding loops, and Metallothionein (Avram et al., 2000). Beyond these zinc motifs, ZFPs encompass several domains that fulfill varied roles in cell biological processes. Notable domains include BTB (Broad-Complex, Tramtrack, and Bric-a-brac), the Krüppel-Associated Box (KRAB) domain, SET domain, and SCAN (SRE-ZBP, CTfin51, AW-1, and Number 18 cDNA) domain(Dunham et al., 2012; Farh et al., 2015; Frontini et al., 2002). The diversity inherent in zinc finger motifs and associated domains endows ZFPs with the ability to assume diverse roles in gene regulation across various cellular environments and in response to various stimuli(Dolfini, Zambelli, Pedrazzoli, Mantovani, & Pavesi, 2016).

BCL11A possesses multiple domains that have the potential to carry out specific functions, such as six C2H2 zinc fingers, one C2HC zinc finger, a NuRD-interacting domain, an acidic domain, and a proline-rich domain (Martyn et al., 2018). BCL11A is expressed in different splice variants and interacts with numerous protein partners, but it is not yet clear which domains and partners are essential for its function(Jawaid, Wahlberg, Thein, & Best, 2010). Further investigation is needed to define the critical domains and protein partners that contribute to BCL11A’s functionality. To systemically define BCL11A functional domains for globin gene repression we discussed in detail the structure and function of zinc finger domains in BCL11A and their implications for the regulation of gamma-globin gene expression. The study investigated the role of two zinc finger domains, ZNF4 and Znf2, in BCL11A using CRISPR-Cas9-mediated targeted genomic deletions. The study found that Znf4 and Znf5 adopt the canonical C2H2 zinc finger fold and make numerous hydrogen-bonding and van der Waals contacts with DNA strands to contribute to the binding affinity and specificity, while Znf3 and Znf6 contribute less significantly to DNA binding.

The study evaluated the efficacy of BCL11A inhibition using sgRNAs and the effect of Znf4 and Znf2 on gamma-globin expression. The results showed a significant increase in gamma-globin expression in both CD34 and K562 cells when targeting Znf4, while the effect of Znf2 was comparatively low on fetal hemoglobin induction. This suggests that Znf4 plays a critical role in regulating gamma-globin gene expression and may be a potential therapeutic target for hemoglobinopathies. The results presented in this study offer convincing evidence that the repression of HbF in adult cells is primarily achieved through direct repression mediated by the promoter. Therefore, previous models that suggested BCL11A primarily repressed HbF from a distance from the g-globin promoters (Sankaran et al., 2011; Xu et al., 2010) are no longer viable. The process of hemoglobin switching appears to be less complex than previously assumed. Although chromosomal looping between the LCR and downstream globin genes is essential for high-level expression (Deng et al., 2014), the absence of BCL11A shifts LCR interactions to the g-globin gene (Xu et al., 2010), leading us to suspect that the presence of BCL11A at the g-globin promoter may be incompatible with stable loop formation. Nonetheless, it is still possible that BCL11A plays other roles in the b-globin cluster, such as in the vicinity of HBBP1 or the LCR, that have not been ruled out.

## Notes

### Competing Interest Statement

The authors have declared no competing interest.

### Summary of Updates

In this revised submission to bioRxiv, the author list has been updated for accuracy and completeness. The modifications enhance the manuscript's precision and reflect the collaborative contributions of all relevant authors. This version aims to provide a more comprehensive and transparent representation of the research team involved in the study.

